# Human Taste Cells Express ACE2: a Portal for SARS-CoV-2 Infection

**DOI:** 10.1101/2021.04.21.440680

**Authors:** Máire E Doyle, Ashley Appleton, Qing-Rong Liu, Qin Yao, Caio Henrique Mazucanti, Josephine M Egan

**Author notes:** Address Correspondence to: Josephine M Egan, M.D., National Institute on Aging/Intramural Program, 251 Bayview Blvd, Baltimore MD 21224, and Máire E Doyle, Ph.D. Funded by the NIA/NIH intramural research program. Project Number 1ZIAAG000291-13, JME. The authors declare that there are no conflicts of interest.

## Abstract

Loss and changes in taste and smell are well-reported symptoms of SARS-CoV-2 infection. The virus targets cells for entry by high affinity binding of its spike protein to cell-surface angiotensin-converting enzyme-2 (ACE2). It was not known whether ACE2 is expressed on taste receptor cells (TRCs) nor if TRCs are infected directly. Using an *in-situ* hybridization (ISH) probe and an antibody specific to ACE2, it seems evident that ACE2 is present on a subpopulation of specialized TRCs, namely, PLCβ_2_ positive, Type II cells in taste buds in taste papillae. Fungiform papillae (FP) of a SARS-CoV-2+ patient exhibiting symptoms of COVID-19, including taste changes, were biopsied. Based on ISH, replicating SARS-CoV-2 was present in Type II cells of this patient. Therefore, taste Type II cells provide a portal for viral entry that predicts vulnerabilities to SARS-CoV-2 in the oral cavity. The continuity and cell turnover of the FP taste stem cell layer of the patient were disrupted during infection and had not fully recovered 6 weeks post symptom onset. Another patient suffering post-COVID-19 taste disturbances also had disrupted stem cells. These results indicate that a COVID-19 patient who experienced taste changes had replicating virus in their taste buds and that SARS-CoV-2 infection results in deficient stem cell turnover needed for differentiation into TRCs.

## Introduction

As many as 80% of people infected with SARS-CoV-2 report taste and smell changes, as well as changes in overall oral sensitivity to commonly used condiments and spices. The constellation of sensory symptoms can precede systemic symptoms and therefore has predictive value.^1, 2^ Interestingly, RNA for SARS-CoV-2 was detected in the submandibular glands and tumor of a patient undergoing tongue surgery for squamous carcinoma two days before the patient developed symptoms and had a positive nasal swab for the virus:^3^ however, taste tissue was not examined. The sensory symptoms need not be unitary as the virus can independently target all or any combination of the senses^4, 5^ and may even cause deficits or changes in a specific taste quality. Branches of five cranial nerves (CN I [smell], V [chemesthesis: heat, pungency], and VII, IX and X [taste, Figure 1A] are involved in relaying those specific senses to the central nervous system. Taste is first discriminated in taste receptor cells (TRCs) within taste buds, which are autonomous organs mostly located in circumvallate (CV), foliate (FL), and fungiform (F) papillae (P) in the tongue. Each taste bud is supplied by nerve fibers, a capillary artery and vein. Solitary taste buds are also buried in the epithelial layer of the uvula, epiglottis, larynx, upper airways and proximal esophagus. TRCs contain the machinery for discriminating five prototypic tastes that can be appreciated from commonly consumed foods: salty (pretzels), sweet (chocolate, sugar, artificial sweeteners), bitter (coffee), umami (monosodium glutamate, sweet amino acids) and sour (citrus); one, more than one or all may be altered (dysgeusia) or absent (ageusia) due to SARS-CoV-2 infection. Taste buds contain just three distinct types of TRCs for mediating taste discrimination: Type I (salty), Type II (sweet, bitter and umami; these are discriminated on the basis of specific receptor engagement on the cell surface while sharing downstream signal transduction mechanisms) and Type III (sour) (Figure 1A).^6, 7^ Solitary chemosensory cells containing similar bitter receptors and signal transduction machinery as are present in Type II cells reside in trachea, extrapulmonary bronchi and in the lung hilus, where based on rodent investigations they ‘sense’ the microenvironment and regulate respiration rate through CN X; bitter stimuli and cholinergic activation lead to decreased respiratory rates.^8^ All taste buds have the same three TRC types, though ratios of cell types vary (Figure 1A). For example, Type I cells are rare in FP.^9^ These TRCs are not neurons: they are modified epithelial cells. ACE2 expression has not been found in the neurons of the geniculate ganglion^10, 11^ whose fibers innervate the FP *via* CN VII, or in sensory afferents of CN X^12^ that innervate the solitary buds of the epiglottis, larynx, and chemosensory cells of the airways. Additionally, CN V neurons, which mediate chemesthesis, do not express ACE2.^13, 14^ It therefore seemed unlikely that alterations in taste and chemesthesis are due to SARS-CoV-2 invasion of sensory afferents of four CNs, but more likely due to direct infection of the taste buds, lingual epithelium and oral mucosa. Epithelial cells shed into saliva, the source of which could be lingual epithelium, salivary glands and oral mucosal surfaces, from SARS-CoV-2 infected patients were recently reported to contain low ACE2 expression, and in some cases, viral RNA.^15^ TRCs or chemosensory cells were not studied in that report.

**Figure 1.**
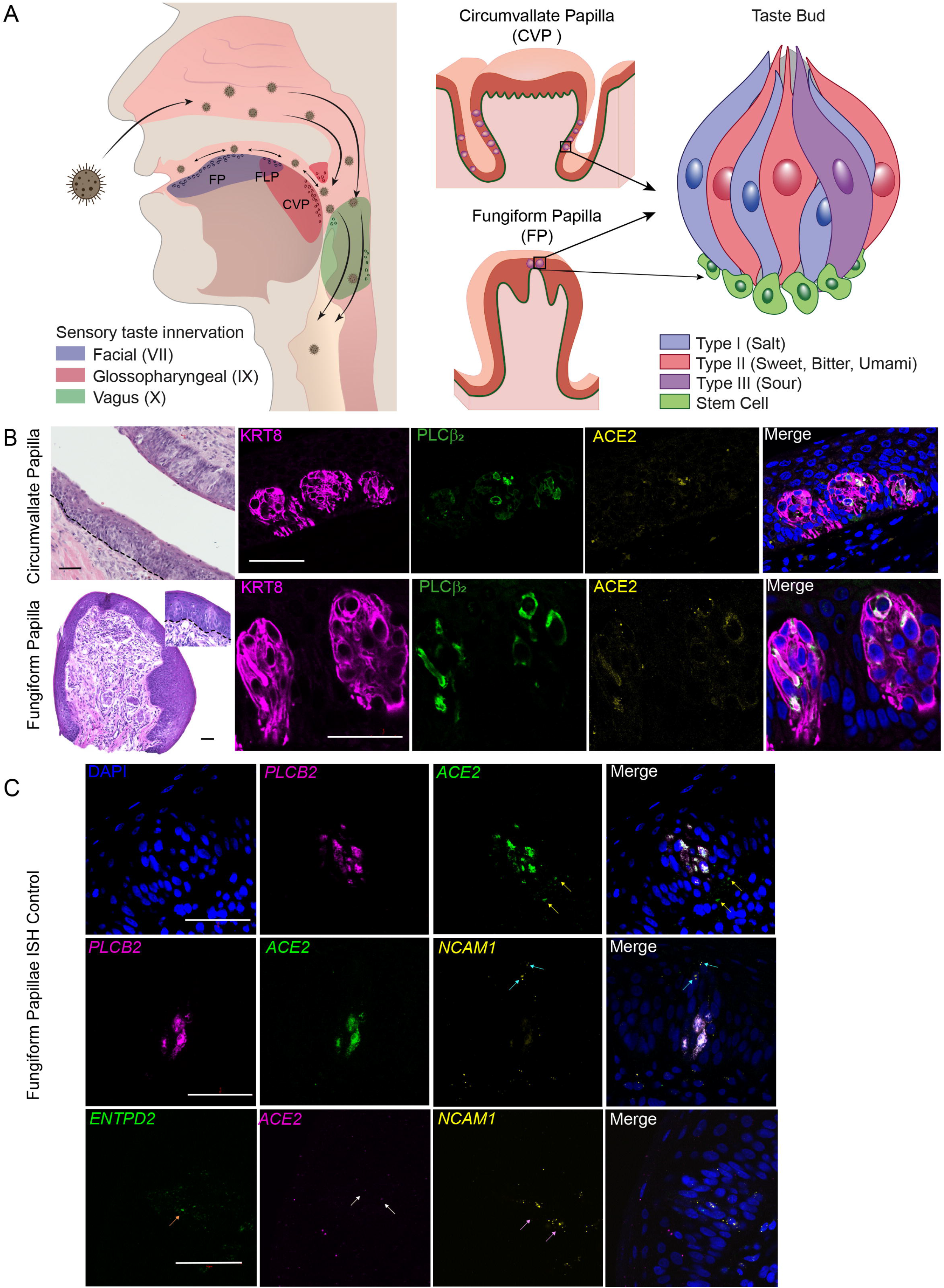
The receptor for SARS-CoV-2 angiotensin converting enzyme 2 (ACE2) is on Type II taste bud cells in taste papillae of the tongue. Panel A shows the distribution of taste buds and chemosensory cells in the oropharyngeal cavity and how inhaled virus may infect the tongue and oropharyngeal areas. Branches of three cranial nerves (CN VII, IX and X) are involved in relaying taste information to the central nervous system. Taste is first discriminated in taste receptor cells (TRCs) within taste buds located in circumvallate (CVP), foliate (FLP) and fungiform papillae (FP) in the tongue. Three defined TRCs relay five prototypic tastes. Stem cells immediately surrounding the taste bud receive signals from taste cells prompting differentiation into a replacement TRC. Circles on tongue, uvula, epiglottis and oropharyngeal areas represent taste buds and chemosensory cells. Panel B, top row shows hematoxylin and eosin (H&E) and immunofluorescent staining (IFS) of CVP (post-mortem) with taste buds embedded in the epithelial layer. Keratin 8 (KRT8) is a cytoskeletal marker of all TRCs while phospholipase C beta 2 (PLCβ_2_) is an obligatory signal molecule in all Type II cells. ACE2 and PLCβ_2_ were colocalized (merged signals) in IFS images. Nuclei are shown in blue stained with 4′,6-diamidino-2-phenylindole dihydrochloride (DAPI). Likewise, H&E staining of a fresh FP with two taste buds (insert), and IFS for KRT8, PLCβ_2_ and ACE2 shows co-localization of the latter two proteins. Dashed lines in H&E of CVP and FP indicate the location of the line of stem cells. Panel C shows *in situ* hybridization (ISH) images of FP. Top panel, probes for *PLCB2* and *ACE2* confirm their co-localization in a fresh FP taste bud, nuclei shown in blue. Note the yellow arrows indicate two areas outside the taste bud where *ACE2* signal is found in the absence of *PLCB2*. Middle panel, co-localization of *ACE2* and *PLCB2* in the same cell is observed and there is no overlap of the Type III cell marker neural cell adhesion molecule 1 (*NCAM1*, light blue arrows),^7^ with either of these two markers. Likewise, bottom panel, shows no overlap of *ACE2* (taste cell positivity indicated by two white arrows) with the probe for the transcript of the Type I cell marker ectonucleoside triphosphate diphosphohydrolase 2 (*ENTPD2*, orange arrow)^7^ and the Type III marker *NCAM1* (taste cell positivity indicated by two pink arrows). Scale bars = 50µm.

## Materials and Methods

### Study design, study population and setting

Human circumvallate papillae (CVP) tissue was obtained from cadaveric tongues and placed in formalin (National Disease Research Interchange) until processing at the NIA. Fresh human fungiform papillae (FP), ≤ eight per participant, were obtained with IRB approval (IRB/NIH # 2018-AG-N010, -N322) and with participants’ written consent. All biopsies were carried out in the IRP NIA Clinical Research Unit, Baltimore, MD. FP were excised after topical application of 1% lidocaine using sterile curved spring micro-scissors (McPherson-Vannas, WPI, Sarasota, FL) Type # SR5603 (Roboz Surgical Instrument Co, Gaithersburg, MD). Individual papillae to be used for immunohistochemistry or ISH were immediately placed in 4% paraformaldehyde (PFA, Fisher Scientific, Atlanta, GA), cryoprotected with 20% sucrose (Millipore Sigma, St. Louis, MO) overnight at 4°C and frozen in optimal cutting temperature media (Tissue Tek O.C.T. Compound, Sakura Fintek, St. Torrance, CA) and stored at -80°C until use.

### Immunostaining of human lingual tissue

CVP tissue and FP were cryosectioned (10 µm thick) using a Leica CM 1950 cryostat, mounted onto ColorFrost Plus Micro slides (Fisher Scientific) and then stored at -80°C. Immunostaining was performed as described previously.^9^ To permeabilize the cells in the tissue, slides were placed in Tris-buffered saline (TBS) (pH, 7.4; Quality Biologicals, Gaithersburg, MD) with 0.2% Triton-X 100 (Millipore Sigma) for 5 minutes at room temperature. They were then washed three times (2 minutes) in TBS. Antigen retrieval was performed by placing the slides in 10 mM of sodium citrate buffer (pH, 6.0; Vector Laboratories, Burlingame, CA) at 95°C for 30 minutes. The slides were left to cool at room temperature in the citrate buffer for a further 30 minutes and were then rinsed in water and then TBS as before. Sections were incubated with normal goat serum block consisting of 2% goat serum, 1% OmniPur® BSA Fraction V (Millipore Sigma), 0.1% gelatin, (Millipore Sigma), 0.1% Triton X-100 (Millipore Sigma), 0.05% Tween 20 (Millipore Sigma), and 0.05% sodium azide (Millipore Sigma) in TBS for 1 hour at room temperature. Sections were then incubated with primary antibodies (Table 1) diluted in the same normal goat serum block at 4°C overnight. Tissue sections were rinsed with TBS with 5% tween (TBST) and incubated for 1 hour with fluorescently labelled secondary antibodies (Table 1) at 1:1000 dilution, then washed with TBST. After nuclear staining (4′,6-diamidino-2-phenylindole dihydrochloride, DAPI, Sigma Aldrich) sections were mounted with Fluoromount G (Southern Biotechnology, Birmingham, AL). Controls used were incubation without primary antibody, isotype controls, and pre-absorption with a 5-fold excess of the blocking peptide (ACE2). Appropriate no-primary-antibody controls were prepared with each individual batch of slides. Confocal fluorescence images were captured using Zen Black Version 14 software on a Zeiss LSM-880 (Oberlochen, Germany) confocal microscope, brightness and contrast were adjusted globally. All figures were compiled in Adobe Illustrator 2021 (San Jose, CA)

**Table 1.**
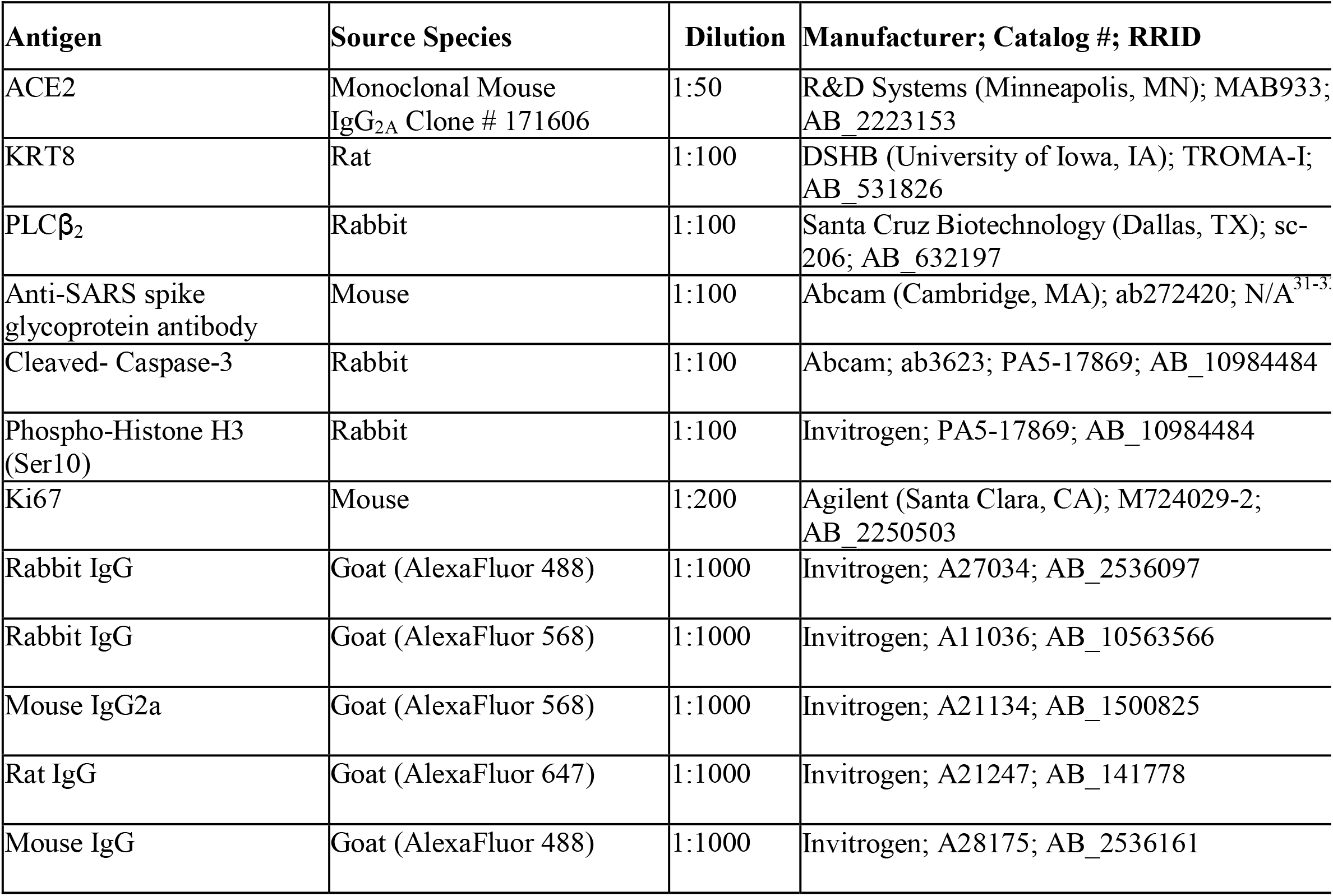
Primary and secondary antibodies used, their dilutions and RRID numbers.

### RNAscope *in situ* hybridization (ISH)

RNAscope probes were all from Advanced Cell Diagnostics (Newark, CA; Table 2). V-nCoV2019-S (Cat# 848561) is an antisense probe specific to the viral genomic positive strand RNA of the spike protein sequence and V-nCoV2019-orf1ab (Cat# 859151) is a sense probe specific to the viral ORF1ab (Open Reading Frame 1ab) negative strand RNA produced when the virus is replicating^16^. Taste receptor cell marker probe for Type I cell was *ENTPD2* (Cat# 507941), for Type II cell was *PCLB2* (custom designed), and Type III cell was *NCAM1* (Cat# 421468). *ENTPD2* is translated into ectonucleoside triphosphate diphosphohydrolase 2 which is expressed only on Type I cells,^17^ while *NCAM1* is translated to a neural cell adhesion molecule that is a cell surface marker for Type III cells.^18^ FP were sectioned at 10 µm using RNAse-free conditions on a Leica CM 1950 cryostat, mounted onto ColorFrost Plus Micro (Fisher Scientific) slides and stored at -80°C in a slide box that was placed in a sealed Zip-loc™ bag until use. Retrieving mRNA targets from the fixed frozen FP sections, pretreatment, probe hybridizations, and labeling were all performed according to RNAscope Multiplex Fluorescent Detection Kit v2 (Cat# 323100) protocol: the negative control probe was a universal control probe targeting the bacterial *Dapb* gene (GenBank accession number: EF191515) from the Bacillus subtilis strain (Cat# 320871). Positive controls were probes to the human transcripts *POLR2A* (C1) and *PPIB* (C2), and *UBC* (C3) (Cat# 320861) for RNAscope Multiplex Fluorescent Assay. Images were acquired using Zen software on a Zeiss LSM 880 confocal microscope.

**Table 2:**
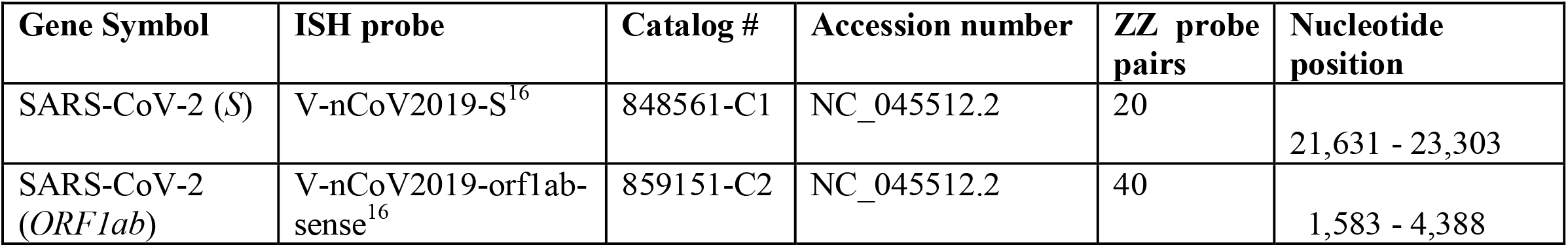
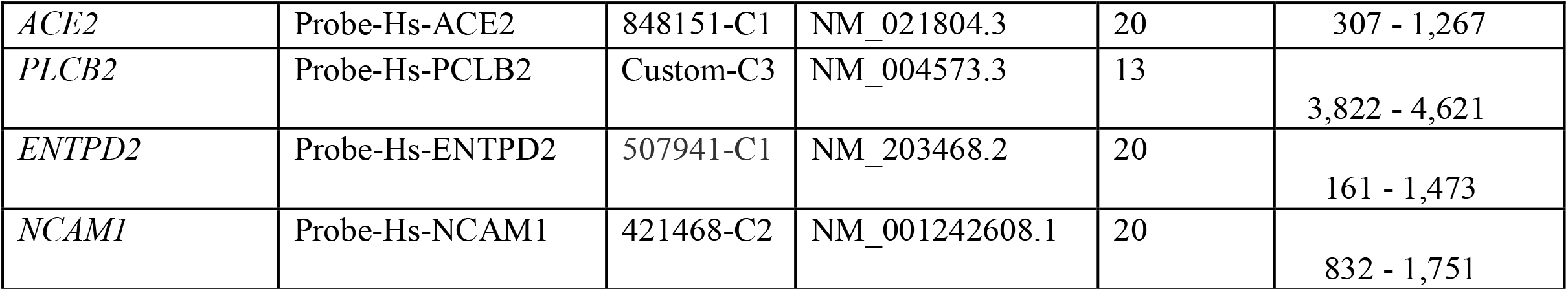
RNAscope ISH probes. All probes were used were off the shelf from Advanced Cell Diagnostics with the exception of PLCB2 which was designed in house by the authors. The website used to obtain the NM accession numbers was https://www.ncbi.nlm.nih.gov/nuccore

| Gene Symbol | ISH probe | Catalog # | Accession number | ZZ probe<br>pairs | Nucleotide<br>position |
| --- | --- | --- | --- | --- | --- |
| SARS-CoV-2 (*S*) | V-nCoV2019-S<sup>16</sup> | 848561-C1 | NC_045512.2 | 20 | 21,631 - 23,303 |
| SARS-CoV-2<br>(*ORF1ab*) | V-nCoV2019-orf1ab-<br>sense<sup>16</sup> | 859151-C2 | NC_045512.2 | 40 | 1,583 - 4,388 |
| *ACE2* | Probe-Hs-ACE2 | 848151-C1 | NM_021804.3 | 20 | 307 - 1,267 |
| *PLCB2* | Probe-Hs-PLCB2 | Custom-C3 | NM_004573.3 | 13 | 3,822 - 4,621 |
| *ENTPD2* | Probe-Hs-ENTPD2 | 507941-C1 | NM_203468.2 | 20 | 161 - 1,473 |
| *NCAM1* | Probe-Hs-NCAM1 | 421468-C2 | NM_001242608.1 | 20 | 832 - 1,751 |

### Quantification of Percentages of Proliferating Cells

The numbers of cells positive for Ki67 and phosphorylated histone H3 (PHH3) were determined using the Cytonuclear FL algorithm in the HALO® image analysis software platform (HALO platform 2.2, Indica Labs, Albuquerque, NM) that identified Ki67-positive nuclei (green), PHH3-positive nuclei (red), and DAPI stained nuclei (blue) on immunostained sections imaged on the LSM 880 using the 20X objective. The percentage of Ki67 positive cells was determined as a percentage of the total number of DAPI positive nuclei. The percentage of Ki67 positive cells positive for the mitotic marker PHH3 was also determined. Cells in three to four sections from each of 2 FPs per individual were counted. One-way analysis of variance (ANOVA) with Tukey’s multiple comparison was performed using GraphPad Prism 9 (GraphPad Software Inc., San Diego, CA). The data are presented as mean ±SEM and p < 0.05 is considered statistically significant.

## Results

Humans have approximately 5,000-10,000 taste buds, of which about half are buried on the sides of CVP (Figure 1A, B).^19^ The presence of ACE2 in taste buds within CVP of post-mortem tissue was investigated by immunofluorescent staining (IFS) and found it to be co-expressed with phospholipase C β_2_ (PLCβ_2_), a marker and obligatory signal transduction molecule in Type II TRCs (Figure 1B, top panel) ^20^. Taste buds within FLPs, which are vertical folds on the sides of the tongue, were not sufficiently preserved for definitive ACE2 IFS.

Obtaining fresh FP, on the other hand, is a relatively simple biopsy procedure. There are 150-200 FP on the front half of the tongue and each FP contains zero to two buds maximum on its surface (Figure 1B, bottom panel, for example). TRCs are replaced approximately every 14 days from replicating stem cells underneath the epithelial layer of FP.^21^ ACE2 was found to be present in TRCs within taste buds of freshly biopsied FP, where again it colocalized with PLCβ_2_ by IFS (Figure 1B, bottom panel). This was confirmed by *in situ* hybridization (ISH; Figure 1C, top panel and middle panel), note the presence of *ACE2* in the area outside of the taste bud. It therefore seems evident that taste buds provide a portal for SARS-CoV-2 entry. *ACE2* (note white arrows) was not found to be co-expressed with Types I and III cell markers (ISH; Figure 1C, bottom two panels).

Participant #114 (female with controlled hypertension, 45yrs old) in the study contracted SARS-CoV-2 (SARS-CoV-2^+^ by PCR) and she reported changes in sweet taste (chocolate did not taste like chocolate) and in chemesthesis (a curry meal tasted ‘white’) beginning 15 days previously. Her tongue had enlarged, red-appearing FP, contrasting with the shiny condition of her FP 3 months later (Figure 2A). Four FP were removed for histology. Of these four FP, just one contained taste buds (Figure 2B). An RNAscope antisense probe specific to the genomic positive strand RNA (for proof of viral infection) of the spike protein (*S*) sequence of SARS-CoV-2 and a sense probe specific to the *ORF1ab* negative strand RNA (for proof of viral replication) indicated the presence of replicating SARS-CoV-2 in *PLCB2* positive cells (participant #114; Figure 2C). Note the arrow pointing to another viral positive cell in the neighboring taste bud. A sex- and age-matched control FP was negative for the virus (Supplemental Figure 1A). The virus was present in lamina propria of participant #114 both by ISH and IFS (Supplemental Figure 1B, C respectively). The stem cell layer within the FP of #114 had disruptions based on immunostaining for Ki67 (a marker of cell turnover): this had improved on a subsequent biopsy 6 weeks later by which time her sense of taste had recovered (Figure 2D, left-hand side images). An age and sex matched control was used as a healthy control for #114. Participant #089 (male with no pre-existing medical conditions, 63yrs old) first donated FP in 2019 (pre-SARS-CoV-2). He was biopsied 6 weeks after being SARS-CoV-2^+^ by PCR but at a time when he still had mild dysgeusia specifically his sweet (chocolate was almost tasteless) and bitter (coffee tasted like mud) sensations were still impaired by history. At this time, his other viral symptoms namely chills, muscle aches, joint pain, brain ‘fog’, had abated and he had returned to work. He was biopsied again at 10 weeks post SARS-CoV-2. No virus was present in his FP (Supplemental Figure 1D): however, his stem cell layer had disruptions 6 weeks post infection compared to 2019 (#089 Pre), and the disruptions were less obvious and taste perception was recovering (coffee was now perceived as tasteless, but not muddy) at 10 weeks (Figure 1D, right hand side images). The total number of cells positive for Ki67 (as a percentage of 4′,6-diamidino-2-phenylindole DAPI-positive nuclei) and the percentage of Ki67 cells positive for the mitotic marker phosphorylated histone H3 (PHH3) in 3-4 sections from each of two FP from participants at the three timepoints shown in Figure 2D were counted (Figure 2E). One-way ANOVA showed that the percentage of Ki67 positive cells in FP of #114 (during) was significantly lower than control and 6 weeks post-COVID-19 (F_(2,21)_=25.2, p<0.0001). The percentage values of Ki67 positive cells in FP of participant #089 at 6 and 10 weeks post-COVID-19 were significantly lower than those of his pre-COVID-19 value (F_(2,18)_=6.97, p=0.0057). One-way ANOVA also showed the percentage of PHH3/Ki67 positive cells in the FP sections from #114 during the course of COVID-19 was significantly lower than those in sections taken from the healthy, sex and aged matched control and #114 6 weeks post-COVID-19 (F_(2,21)_=19.35, p<0.0001). The percentage of PHH3/Ki67 positive cells in FP sections from #089 at 6 and 10 weeks post-COVID-19 were significantly lower than those of the pre-COVID-19 FP sample (F_(2,18)_=21.23, p<0.0001). A marker of cellular apoptosis, cleaved caspase 3, was not expressed in the stem cell layer during COVID-19 in participant #114 or during the period of taste dysfunction of participant #089 as shown in Figure 2F.

**Figure 2.**
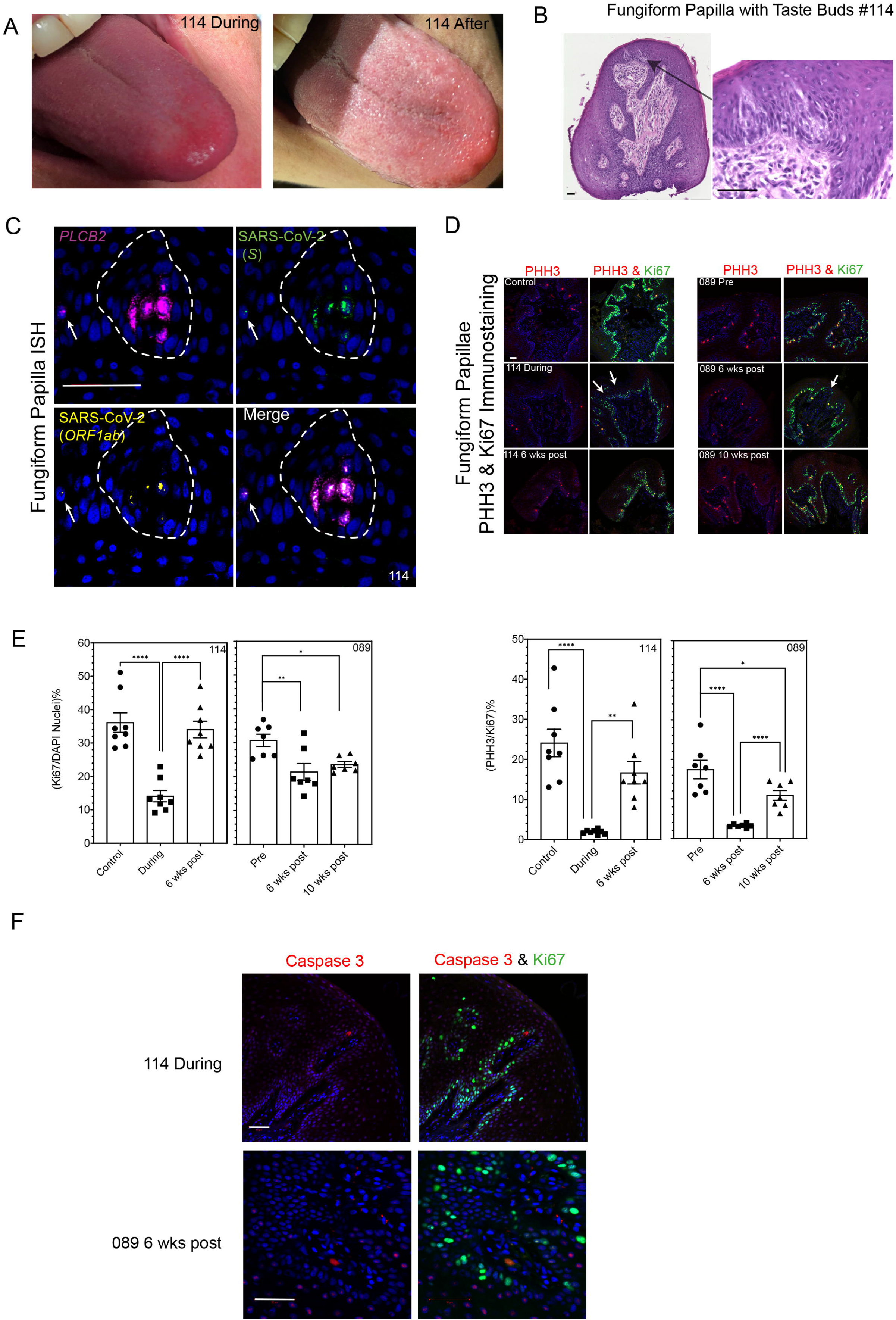
Evidence of SARS-CoV-2 in human FP. Panel A shows tongue photographs of patient #114 moments prior to biopsy of her FP during the course of COVID-19 and 3 months later. Panel B shows H&E staining of a section through the FP from patient #114 that contained two taste buds; the consecutive section was used for ISH outlining the presence of viral particles (SARS-CoV-2 *S*) probe for the Spike mRNA and the SARS-CoV-2 (*ORF1ab*) probe for the replicating virus, shown in Figure 2C. Panel C, an antisense probe specific to the genomic positive strand RNA of the spike protein (*S*) sequence of SARS-CoV-2 and a sense probe to the SARS-CoV-2 *ORF1ab* negative strand RNA indicate the presence of replicating virus in *PLCB2* positive cells (participant #114). Note the arrow pointing to another viral positive cell in the neighboring taste bud. Panel D demonstrates the proliferation of the stem cell layer of the FP by immunostaining for the marker of all active phases of cell cycle Ki67 and the late G2 and M phase markers phosphorylated histone H3 (PHH3). Top left-sided images show a FP from an age- and sex-matched control participant for #114 compared with during and post SARS-CoV-2 infection. White arrows indicate the breaks in the otherwise continuous layer of stem cells. Right-sided panels show a continuous stem cell layer in participant #089 pre, but multiple breaks especially at 6 weeks, and less at 10 weeks, post SARS-CoV-2. Panel E shows the percentages of total cells (as determined by DAPI stained nuclei) positive for Ki67 and the percentage of Ki67 positive cells that are also positive for PHH3 (mean ± SEM, **** = p<0.0001, ** = p <0.01, * = p < 0.05). Panel F shows that the proliferating taste stem cells do not express cleaved Caspase 3. Scale bars = 50µm.

## Discussion

We discovered that oral infection and replication with SARS-CoV-2 occurred in taste bud Type II cells. Virus may be transported to the oral cavity on air eddies containing virus in droplets directly into mouth and/or in droplets, nasal mucous or epithelial cells shed from the nose (Figure 1A). While SARS-CoV-2 infection is noted in oral cavity and saliva, no one had shown evidence of its replication in salivary glands, gingiva or oral mucosa, and no one has yet published on taste tissue. Furthermore, previous studies^15, 22, 23^ did not address the cellular location of ACE2 within human taste buds using the highly specific techniques of ISH (mRNA) and IFS (protein). Taste buds do not contain a resident immune system^24^ and TRCs account for all cells within taste buds. In contrast to SARS-CoV-2 infection in the olfactory system^25^ where it is the sustentacular cells that are affected our data show evidence for infection in the taste apparatus *per se i*.*e*. the taste buds. Replication of virus can likely then occur undisturbed and allow for transmission from the taste bud into circulation, and locally infect lingual and salivary gland epithelium, oral mucosa and larynx and even on into the lungs. TRCs within taste buds contain interferon receptors, and a systemic response to virus should eliminate the virus while simultaneously leading to taste changes.^24^ While we did not investigate it directly it is possible that there could be indirect effects on the neuronal^26^ or blood supply to the taste buds as is noted by the detection of ACE2 in the proximity of the taste buds. We further propose that deficient stem cell turnover will result in TRCs not being effectively replaced and that this explains why some people have slow recovery of their complete taste repertoire. The detection of actively replicating SARS-CoV-2 in *PLCB2*-containing cells indicates that the virus has a direct route into TRCs via ACE2 on the type II cells. The presence of ACE2 on Type II cells specifically is very intriguing because these are the cells that ‘taste’ amino acids and ACE2 in the gut has been shown to dimerize with amino acid transporters where it plays a vital role in efficient amino acid absorption.^27^ We hypothesize that ACE2 is also on the chemosensory cells present in airways as their cellular machinery is similar to Type II cells in taste buds. They contain bitter receptors activated by ligands such cycloheximide, a bitter receptor agonist, and require PLCβ_2_ for signal transduction.^8^ Therefore, viral infection of those cells might directly decrease respiratory drive, resulting in worsening oxygen desaturation. The role of chemosensory cells in the upper airways or of TRCs in taste buds during SARS-CoV-2 infection has not been studied because of practical difficulties: obtaining a fresh CVP where numerous taste buds reside and where they are larger than in FP is not a viable option because CVPs have not been shown to regrow^28^; obtaining FP or especially fresh tissue from airways in sick people has associated medical conundrums; there is a paucity of taste buds within FP as there are at most two taste buds but sometimes just one and occasionally none, even in young people^29^; taste buds may be damaged by virus and not be recognizable, especially to an untrained eye; taste buds are buried in the epithelial layer where they occupy less than 1% of the total mass of a papilla (Figure 1A, B) and therefore are easily missed if the whole papilla is not systematically sectioned; and finally, even when present, a taste bud in a FP is approximately 30 µm in diameter, thereby providing a maximum of 3-4 slides with taste bud cells for in depth investigation.

In conclusion by demonstrating the co-localization of SARS-CoV-2 virus, Type II taste cell marker and the viral receptor ACE2, we show evidence for replication of this virus within taste buds that could account for acute taste changes during active COVID-19. This work also shows that proliferation of the taste stem cells in recovering patients may take weeks to return to their pre-COVID-19 state, providing a hypothesis for more chronic disruption of taste sensation, reports of which are now appearing in the medical literature. It is worthwhile noting that the influenza A virus subtype H3N2 that caused the pandemic of 1968-69 resulted in long-term alternations of taste and smell in some patients. Patients suffering from post-influenza hypo- and dysgeusia many years after their infection had disrupted taste bud architecture with decreased numbers of TRCs and were lacking cilia in the pore region of the taste bud.^30^

## Acknowledgements

All authors contributed equally. We thank all the participants of the study (2018-AG-N010) and all the IRP NIA clinical staff.

## List of Non-Standard Abbreviations

ACE2: Angiotensin-Converting Enzyme-2
CN: Cranial Nerve
COVID-19: Coronavirus Disease 2019
CV: Circumvallate
CVP: Circumvallate Papilla(e)
ddPCR: Droplet Digital Polymerase Chain Reaction
ENTPD2: Ectonucleoside Triphosphate Diphosphohydrolase 2
FL: Foliate Papilla(e)
FP: Fungiform Papilla(e)
ISH: *In-Situ* Hybridization
NCAM1: Neural Cell Adhesion Molecule 1
ORF1ab: Open Reading Frame 1ab
PHH3: Phosphohistone H3
PLCβ_2_: Phospholipase C β 2
SARS-CoV-2: Severe Acute Respiratory 41 Syndrome Coronavirus 2
TMPRSS: Transmembrane Protease, Serine
TRC: Taste Receptor Cell(s)

---

Supplemental Figure 1. Panel A shows RNAscope detection in a taste bud of a fungiform papilla (FP) of the transcript for the type II cell marker *PLCB2* and is the negative non-infected control for the SARS-CoV-2 *ORF1ab* probe in a taste bud biopsied from a non-infected age and sex matched control for #114. Nuclei stained by DAPI, shown in blue. Panel B shows the sense probe targeting the viral *ORF1ab* mRNA demonstrates positivity within the lamina propria of participant #114 during infection. Panel C shows immunostaining in red for the SARS spike protein in the lamina propria of a FP from participant #114 during infection. Panel D there is no evidence of SARS spike protein in the lamina propria of participant #089 post-COVID-19. Scale bars = 50µm.

