## Supplementary figures and images for "Human Taste Cells Express ACE2: a Portal for SARS-CoV-2 Infection"

### Supplemental Figure 1

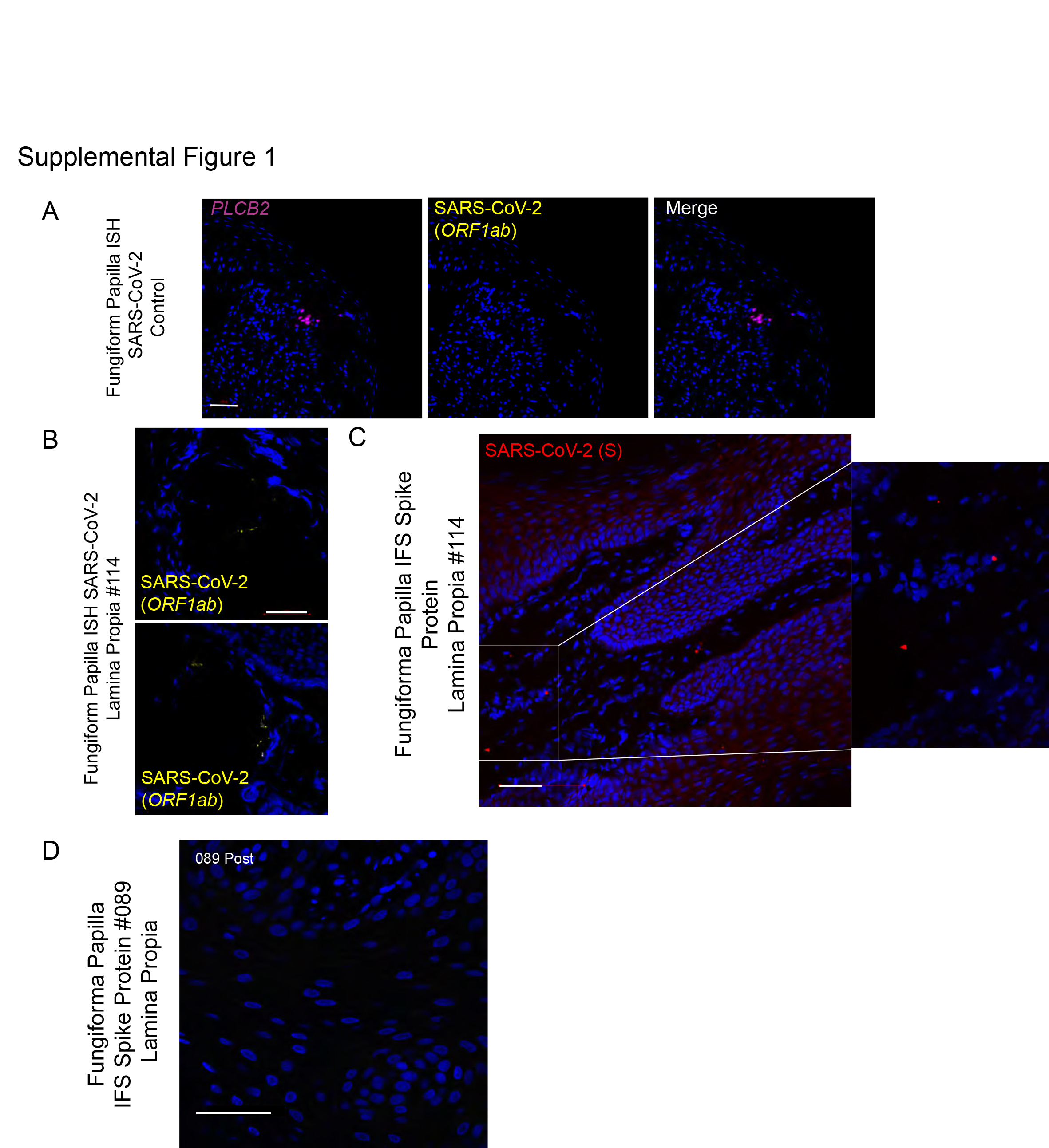
